# New transcriptional circuit evolved by coding sequence changes in a master regulator followed by *cis*-regulatory changes in its target genes

**DOI:** 10.1101/565234

**Authors:** Candace S. Britton, Trevor R. Sorrells, Alexander D. Johnson

## Abstract

While changes in both the coding-sequence of transcriptional regulators and in the *cis*-regulatory sequences recognized by them have been implicated in the evolution of transcriptional circuits, little is known of how they evolve in concert. We describe an evolutionary pathway in fungi where a new transcriptional circuit (**a**-specific gene repression by Matα2) evolved by coding changes in an ancient master regulator, followed millions of years later by *cis*-regulatory sequence changes in the genes of its future regulon. We discerned this order of events by analyzing a group of species in which the coding changes in the regulator are present, but the *cis*-regulatory changes in the target genes are not. In this group we show that the coding changes became necessary for the regulator’s deeply conserved function and were therefore preserved. We propose that the changes first arose without altering the overall function of the regulator (although changing the details of its mechanism) and were later co-opted to “jump start” the formation of the new circuit.

## Introduction

Changes in transcriptional circuits over evolutionary time are an important source of organismal novelty. Such circuits are typically composed of one or more transcriptional regulators (sequence-specific DNA-binding proteins) and their direct target genes, which contain *cis*-regulatory sequences recognized by the regulators. Although much early emphasis was placed on changes of *cis*-regulatory sequences as a source of novelty, it is now clear that coding-changes in the transcriptional regulatory proteins that bind the *cis*-regulatory sequences are also of key importance^1-5^. Indeed, some well-documented changes in transcriptional circuitry require concerted changes in both *cis*-regulatory sequences and the regulators that bind them^6,7^. Although such concerted changes are likely to be widespread, we know little about how they occur.

Here, we study a case in the fungal lineage where gains in *cis*-regulatory sequences in a set of target genes as well as coding changes in the master transcriptional regulator itself are needed for a new circuit to evolve. Specifically, we addressed which came first, the changes in the regulatory protein or the changes in the *cis*-regulatory sequences of its 5-10 target genes. We document that the coding-sequence changes in the regulatory protein occurred millions of years before the gains of the *cis*-regulatory sequences. This observation poses an apparent paradox: how could the protein acquire the needed changes when the new circuit was completed only much later in evolutionary time? We answer this question by revealing a new, unanticipated regulatory scheme present in a small clade of understudied fungal species that readily accounts for this evolutionary trajectory.

## Results

The expression of the cell-type specific genes in fungi provides a powerful system to mechanistically study the evolution of gene expression patterns across a period of several hundred million years^8-15^. The Saccharomycotina clade of fungi, which spans this range of evolutionary time (see Fig. 1), have two types of mating cells, **a** and α, each of which express a set of genes specific to that cell type. Only **a** cells express **a**-specific genes and only α cells express α-specific genes. Both **a** and α cells express an additional set of genes, the haploid-specific genes (Fig. 1a). The proteins encoded by these sets of genes have been intensively studied and are critical for the sexual cycle (mating and meiosis) of these species (see Supplementary Information for more details).

**Figure 1.**
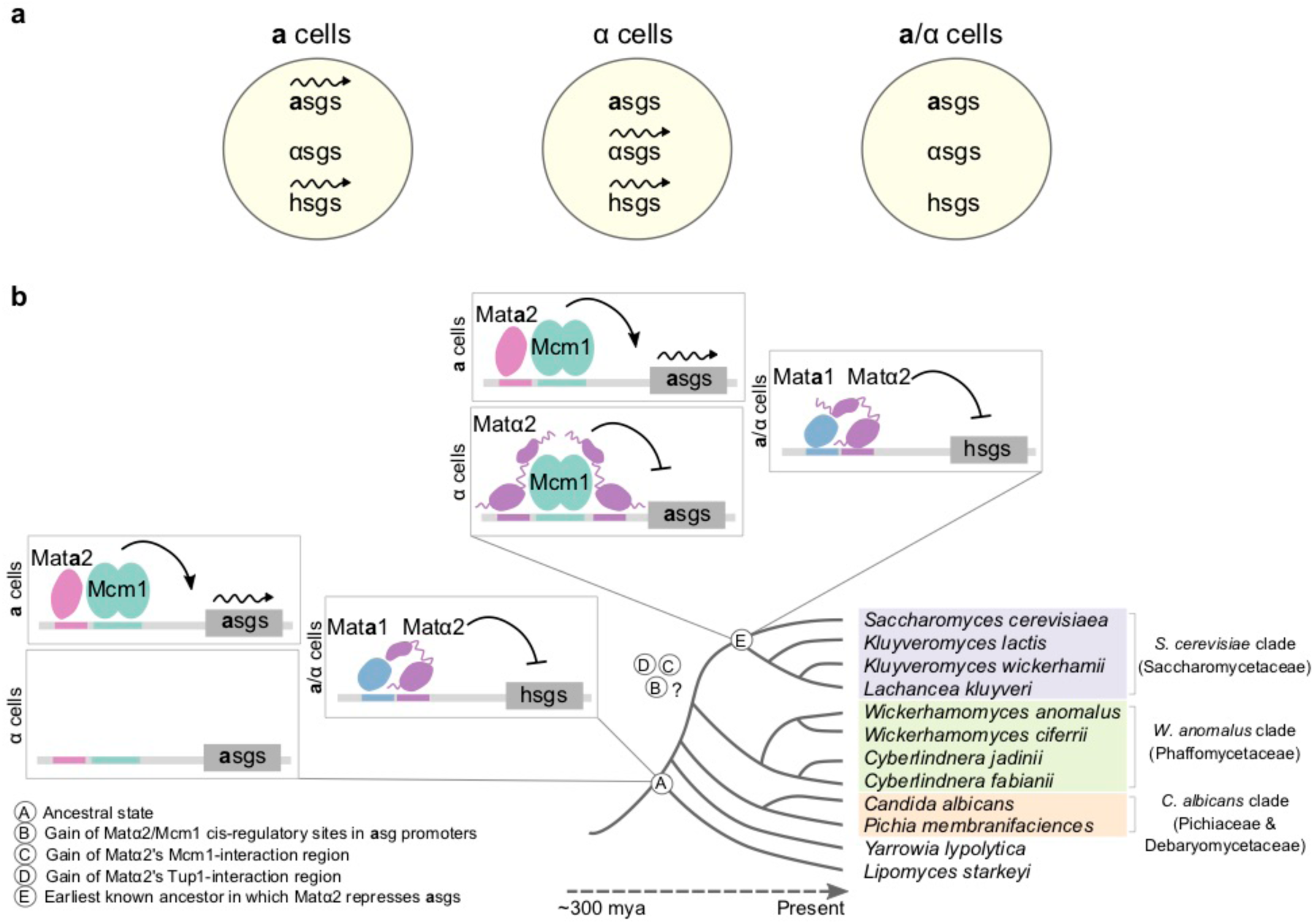
Cell-type specific gene expression in the Saccharomycotina yeast. a. Across the entire Saccharomycotina clade, **a** and α cells each express a set of genes specific to that cell type (**a**-and α-specific genes, or **a**sgs and αsgs, respectively), as well as a shared set of haploid-specific genes (hsgs). **a** and α cells can mate to form **a**/α cells that do not express the **a**-, α-or haploid-specific genes^35^.
b. The mechanism underlying the expression pattern of **a**-specific genes is different among species. In the last common ancestor of the Saccharomycotina yeast (circled A in the figure) transcription of the **a**-specific genes was activated by Mat**a**2, an HMG-domain protein only produced in **a** cells, which binds directly to the regulatory of each **a**-specific gene^9,10^. Much later in evolutionary time (E), repression of the **a**-specific genes by direct binding by Matα2 evolved. Steps B, C, and D must have occurred between steps A and E.

The expression patterns of these three sets of genes are broadly conserved across extant yeasts (as are the genes themselves) and therefore must have been present in the last common ancestor of these species, some hundreds of millions of years ago. However, extant species accomplish the regulation of the cell-type specific genes using different mechanisms (Fig. 1b)^8-10^. For example, in one group of species, the **a**-specific genes are turned on by a transcriptional activator protein (Mat**a**2) that is made only in **a** cells and which binds a *cis*-regulatory sequence upstream of the **a**-specific genes. In another group of species the **a**-specific genes are turned on by a ubiquitous activator and turned off by binding of a repressor (Matα2) that is made only in α and **a**/α cells. Both schemes ensure that the **a**-specific genes are expressed only in **a** cells, although the mechanism behind this pattern is completely different. Previous work established that the activation scheme (activating **a**-specific genes by Mat**a**2) is ancestral to the repression scheme (repression of the **a**-specific genes by Matα2) (Fig. 1b)^9^.

The switch between these two mechanisms of controlling the **a**-specific genes was completed sometime before the divergence of *Saccharomyces cerevisiae* and *Kluyveromyces lactis* (formally known as Saccharomycetaceae, here called the *S. cerevisiae* clade) but after the divergence of this clade and that containing *Candida albicans* and *Pichia membrifaciens* (formally, Pichiaceae and Debaryomecetaceae, here called the *C. albicans* clade) (Fig. 1b). Three events must have occurred for the newer (repression) scheme to have evolved: (1) Matα2 acquired the ability to contact the Tup1-Ssn6 co-repressor, bringing it to DNA to carry out the repression function, (2) Matα2 acquired the ability to bind to DNA cooperatively (through a direct protein-protein contact) with Mcm1, a ubiquitously expressed transcription regulator necessary for **a**-specific gene repression, and (3) the **a**-specific genes (numbering between 5 and 10, depending on the species) acquired *cis*-regulatory sites for Matα2 (Fig. 1b).

To determine the order of these events, we studied Matα2 and the regulation of the **a**-specific genes in a clade that branched from the ancestor before the completion of all three of these events; that is, before the **a**-specific genes had been brought under negative control by Matα2. We reasoned that this group of species might have acquired some, but not all of the changes needed to form the new circuit and therefore might provide clues to its evolutionary history. This approach was made possible by the sequencing of genomes of a monophyletic group of species that branches outside of the last common ancestor of the *S. cerevisiae* clade (formally known as the Phaffomycetaceae) (Fig. 1b)^16,17^. We chose the species *Wickerhamomyces anomalus* as a representative of this branch because it does not undergo mating-type switching, greatly simplifying its analysis, and because we worked out relatively simple procedures to genetically alter it^18^.

The central player in the evolutionary pathway investigated here is Matα2, a homeodomain protein. In *S. cerevisiae*, this protein represses both the **a**-specific genes (in combination with Mcm1, a MADS box protein) and the haploid-specific genes (in combination with Mat**a**1, another homeodomain protein). The latter circuit is ancient, present in all three clades, whereas the former, as discussed above, is a relatively recent acquisition, found only in the *S. cerevisiae* clade (Fig. 1b). The *S. cerevisiae* protein has a modular organization of five functional regions (Fig. 2a). Proceeding from N to C-terminus, these are (1) an N-terminal region that binds the transcriptional co-repressor protein Tup1-Ssn6, (2) a domain involved in heterodimerizing with Mat**a**1 to regulate the haploid-specific genes, (3) an approximately 10 amino acid stretch that binds Mcm1, so that Matα2 and Mcm1 bind cooperatively to the *cis*-regulatory sequences controlling the **a**-specific genes, (4) the homeodomain, required to bind *cis*-regulatory sequences in both the **a**-specific and haploid-specific genes, and (5) a C-terminal region that binds Mat**a**1 and is needed for Mat**a**1 and Matα2 to jointly bind DNA^19-28^. *C. albicans* Matα2, which represents the ancestral state of the Matα2 coding sequence, lacks the region that interacts with Tup1 (region 1) and the residues that interact with Mcm1 (region 3), but shows a significant similarity to *S. cerevisiae* clade Matα2s in the other three regions (Supplementary Fig. 1)^10,23,29^. We previously tested and confirmed these predictions experimentally: when the *C. albicans* Matα2 protein is expressed in *S. cerevisiae*, it cannot repress the *S. cerevisiae* **a**-specific genes^29^. However, if regions 1 and 3 from the *S. cerevisiae* Matα2 are grafted onto the *C. albicans* Matα2, it can now repress these genes^29^.

**Figure 2.**
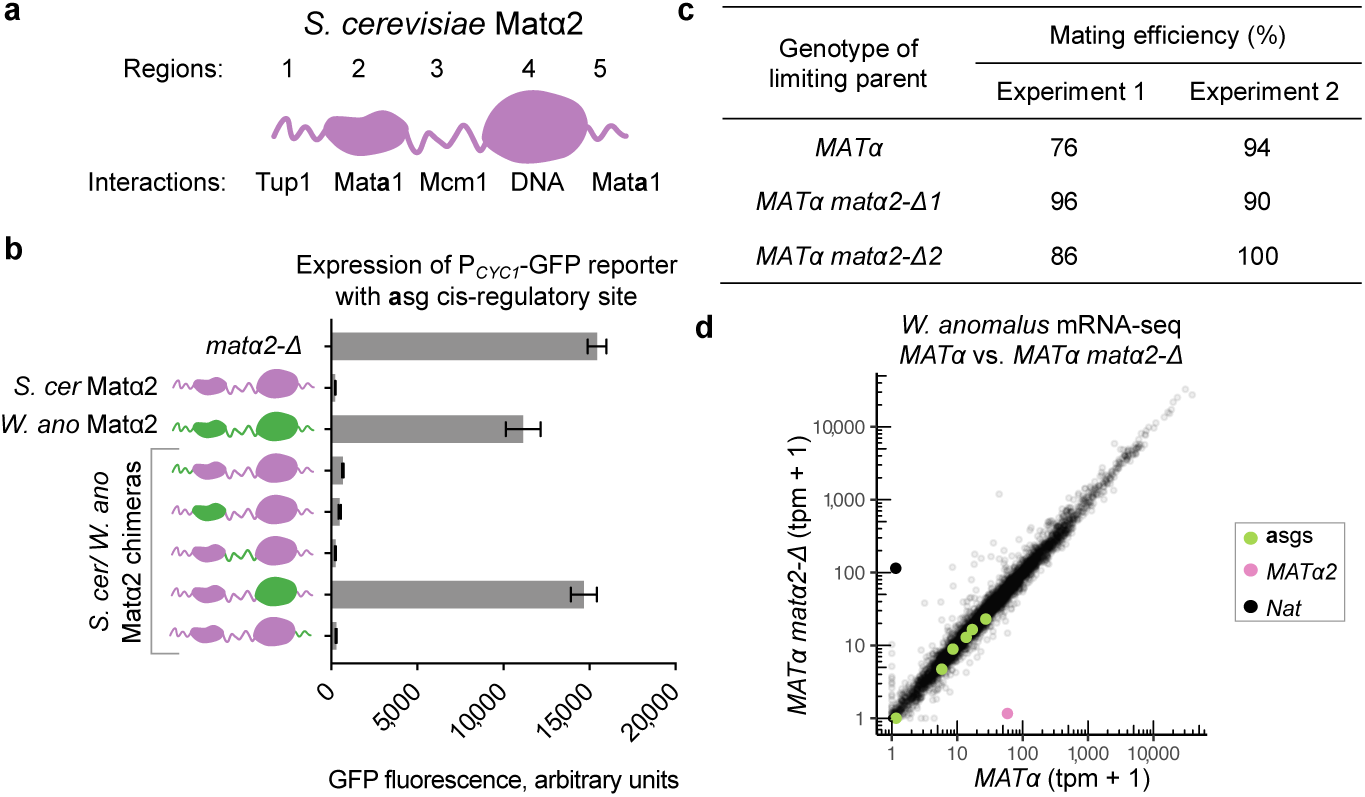
*W. anomalus* Matα2 has functional Tup1 and Mcm1-interacting regions but does not repress the a-specific genes. a. The five modules of the *S. cerevisiae* Matα2 protein. Structural domains are shown as globular, and unstructured regions are shown as wavy lines.
b. Activity of Matα2 derivatives in repressing **a**-specific gene transcription in *S. cerevisiae*. The **a**-specific gene *cis*-regulatory element from the *S. cerevisiae* **a**-specific gene *STE2* gene was placed into a reporter construct and used to measure the ability of *S. cerevisiae* Matα2 (purple), *W. anomalus* Matα2 (green), and hybrid proteins (purple and green) to repress **a**-specific gene transcription. *MATα2* constructs (driven by the *S. cerevisiae MATα2* promoter) were inserted into a *S. cerevisiae* α cell with *MATα2* deleted (*matα2-Δ*). In addition, regions 1-5 of the *W. anomalus* Matα2 (green) sequence were swapped into the *S. cerevisiae* Matα2 (purple) protein and tested for their ability to it. Mean and SD of three independent genetic isolates grown and tested in parallel are shown.
c. In *W. anomalus*, Mat**a**2 is required for **a** cells to mate but Matα2 is not required for α cells to mate. The results of quantitative mating assays (see Supplementary Information) are a. given for wild-type **a** and α cells and various mutant derivatives. The values show the percentage of cells of the limiting parent strain that mated in the experiment.
d. mRNA-seq of wildtype *W. anomalus* α cells (*MATα*) compared to α cells with *MATα2* deleted (*MATα2 matα2-Δ*). *MATα2* expression in transcripts per million (tpm) is shown in pink, confirming it has been deleted. The **a**-specific genes *STE2, AXL1, ASG7, BAR1, STE6*, and *MAT****a****2* are shown in green. The marker used to delete *MATα2* (*Nat*) is shown in opaque black (providing an additional control for the experiment), and all other mRNAs are shown in translucent black. Data from one culture of each genotype is plotted here, and data from independent replicates grown and prepared in parallel and from an independent deletion of *MATα2* are given in Supplementary Fig. 3.
e. **a**-specific gene expression levels in a wildtype *W. anomalus* **a** cells (*MAT****a***) vs. **a** cells with *MAT****a****2* deleted (*MAT****a****2 mat****a****2-Δ*), measured by the NanoString nCounter system^36^. All **a**-specific genes require *MAT****a****2* for wildtype-level expression in the **a** cell. Also shown are expression levels of the α-specific gene *STE3* and the haploid-specific gene *STE4*. Mean and SD of two cultures per genotype, grown and tested in parallel, are shown.

We examined the *W. anomalus* Matα2 protein sequence to determine whether it is more similar to the ancestral (represented by *C. albicans*), or the derived (represented by *S. cerevisiae*) form of Matα2. Alignment of the coding sequence of *W. anomalus* Matα2 with orthologs of Matα2 from many other species indicated that it shares all five functional regions with the *S. cerevisiae* protein (Supplementary Fig. 1). In particular, it has a similar region 1 (the Tup1-interacting region) and a similar region 3 (the Mcm1-interacting region), the regions that the *C. albicans* protein lacks. By swapping these *W. anomalus* regions into the *S. cerevisiae* protein, we confirmed that they are functional in repressing the **a**-specific genes (Fig. 2b). In the course of these experiments, we found, unexpectedly, that the homeodomain of the *W. anomalus* protein contained mutations that prevented its binding to the **a**-specific gene *cis*-regulatory sequence in *S. cerevisiae*, a derived change within this clade alone that we discuss below (Fig. 2b, Supplementary Fig. 1). Similar results were obtained with Matα2 protein from two additional species that branch with *W. anomalus*, indicating that these two conclusions (*W. anomalus* clade Matα2 has functional regions 1 and 3, but cannot bind the *S. cerevisiae* **a**-specific genes) are characteristic of the *W. anomalus* clade rather than a single species (Supplementary Fig. 1d).

The fact that the *W. anomalus* Matα2 protein had acquired the necessary coding changes to interact with Tup1 and Mcm1 but could not bind to the *S. cerevisiae* **a**-specific gene control region raised the question of whether it has a role in regulating the **a**-specific genes in *W. anomalus*. First, we note that the *W. anomalus* **a**-specific genes are indeed cell-type regulated: they are only expressed in **a** cells (Supplementary Fig. 2). In principle, there are three mechanisms that can produce proper expression of these genes: (1) the **a**-specific genes could be under positive regulation by the transcriptional activator Mat**a**2, as is the situation in *C. albicans* and other outgroup species. Alternatively, (2) the **a**-specific genes could be under negative regulation by Matα2 as they are in *S. cerevisiae*, or (3) the **a**-specific genes could be under both positive regulation by Mat**a**2 and negative regulation by Matα2 as seen in *Lachancea kluyveri* and *Kluyveromyces wickerhamii*^29^. We determined that the first possibility is correct. Specifically, we tested the effects of deleting Matα2 from an α cell and Mat**a**2 from an **a** cell on cell-type specific gene expression and mating. We found that Mat**a**2 is required for **a**-specific gene activation and for **a** cells to mate, but that Matα2 does not repress the **a**-specific genes and is not required for α cells to mate (Fig. 2c, d, e, Supplementary Fig. 3). The fact that the *W. anomalus* Matα2 does not regulate the **a**-specific genes can account for the observation that the homeodomain has acquired mutations that compromise its binding to the *S. cerevisiae* **a**-specific genes, as detected in our reporter assays. In addition, we analyzed the *W. anomalus* **a**-specific gene *cis*-regulatory elements bioinformatically and experimentally and found that they possess Mat**a**2-Mcm1, not Matα2-Mcm1 binding sites, entirely supporting the genetic experiments (Supplementary Fig. 4b). These results argue against the possibility that direct **a**-specific gene repression by Matα2 existed in an ancestor of *W. anomalus* but was subsequently lost, as this would have required the independent loss of Matα2 binding sites from all of the **a**-specific genes across numerous species.

The experiments described so far demonstrate that the new circuit—Matα2 repression of the **a**-specific genes—had partially formed by the time the *W. anomalus* clade branched but was not completed until after the divergence of the *S. cerevisiae* clade. The last step in the completion of the circuit was the acquisition of *cis*-regulatory sequences recognized by Matα2 in the **a**-specific genes. This step must have occurred long after Matα2 gained the regions that bind Tup1 and Mcm1. We next address how these changes in the Matα2 protein could have predated the completion of the circuit.

One hypothesis for how Matα2 acquired the two protein-protein interactions needed to repress the **a**-specific genes millions of years before these genes acquired the necessary *cis*-regulatory sequences holds that these interactions are needed for Matα2’s more ancient function in repressing the haploid-specific genes, but only in the *W. anomalus* clade. To test this idea, we analyzed the requirements for haploid-specific gene repression in *W. anomalus*. We deleted *MATα2* and *MAT****a****1* in **a**/α cells and found that they are both necessary for haploid-specific gene repression (Fig. 3a, b, Supplementary Fig. 5). This is the case for virtually all species in the three clades studied to date. However, unlike species outside the clade, we found using genetic experiments that regions 1 and 3 of *W. anomalus* Matα2 (the Tup1-interaction region and the Mcm1-interaction region) are necessary for repression of the haploid-specific genes (Fig. 3c). Finally, we found that a Mcm1 *cis*-regulatory site is also required for the repression of the haploid-specific gene *RME1* (Fig. 3d, e, Supplementary Fig. 6). Taken together with the results of chromatin immunoprecipitation experiments, we show that Matα2, Mat**a**1, and Mcm1 are all required for haploid-specific gene repression in *W. anomalus* and that the portions of Matα2 that interact with Mcm1 and Tup1 are also required. This three-part recognition of the haploid-specific genes in the *W. anomalus* clade was not anticipated from studies of other species. Even in the *S. cerevisiae* clade where Mcm1 and Matα2 are known to interact, this interaction is not required for haploid-specific gene repression. This analysis explains the initially puzzling observation that the key changes in Matα2 that were needed to form the new **a**-specific gene circuit were already in place in the last common ancestor of *S. cerevisiae* and *W. anomalus*, long before the circuit was completed (Fig. 4).

**Figure 3.**
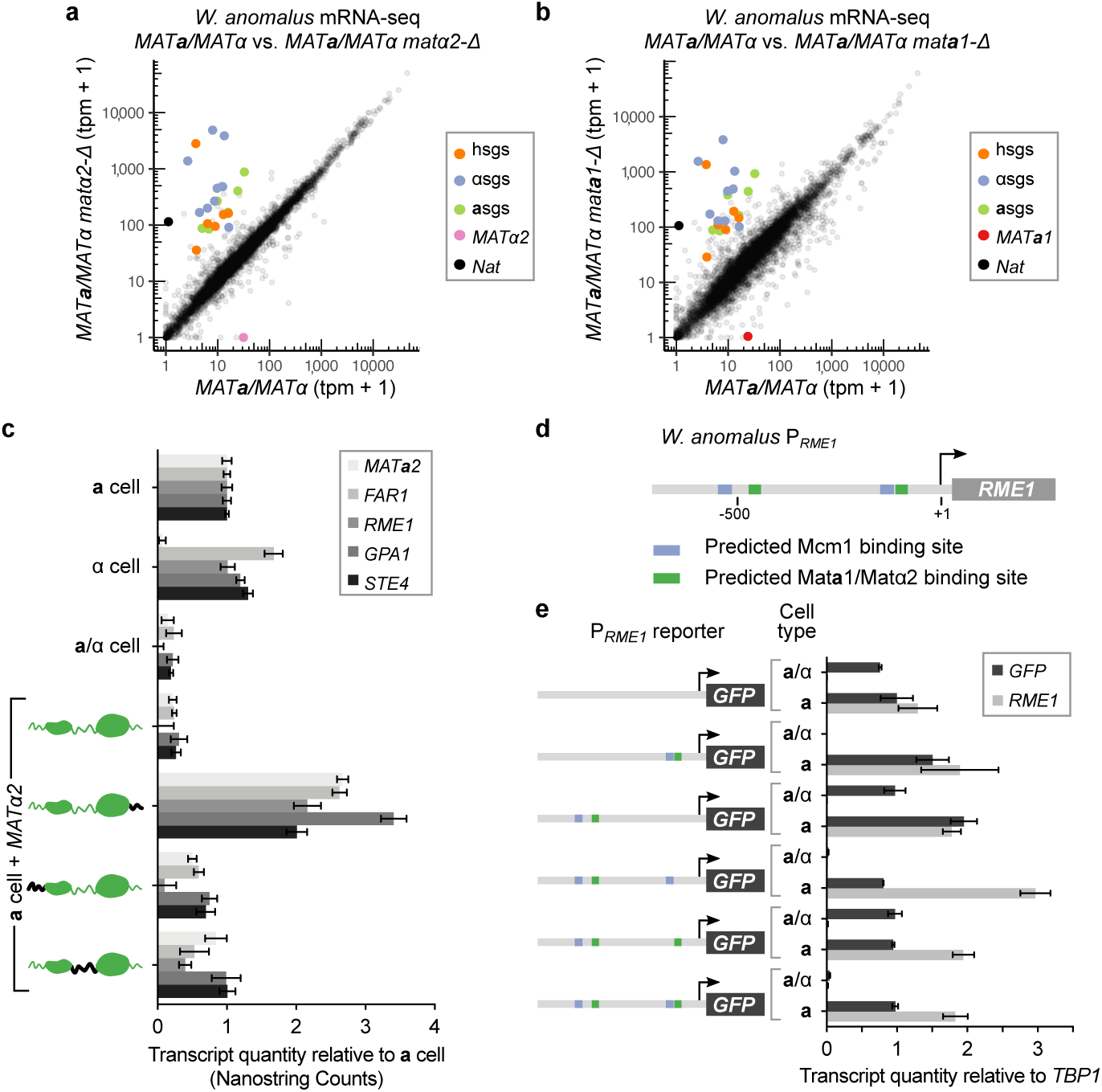
Mata1, Matα2, and a Mcm1 *cis*-regulatory sequence are all required for haploid-specific gene repression in *W. anomalus*. a. Matα2 is required for haploid-specific gene repression. mRNA-seq of wildtype *W. anomalus* **a**/α cell (*MAT****a****/MATα*) compared to an **a**/α cell with *MATα2* deleted (*MAT****a****/MATα matα2-Δ*). As in the previous figure, *MATα2* expression in transcripts per million (tpm) is shown in pink, confirming it has been deleted. The **a**-specific genes *STE2, AXL1, ASG7, BAR1, STE6*, and *MAT****a****2* are shown in green, the haploid-specific genes *STE4, GPA1, FUS3, SST2, RME1*, and *FAR1* are shown in orange, the α-specific genes *SAG1, STE3, STE13, MFα* (multiple copies), *AFB1* (multiple copies), and *MATα1* are shown in blue-gray. The marker used to delete *MATα2* (*Nat*) is shown in opaque black, and all other mRNAs are shown in translucent black. The results show that the **a**-, α-and a. haploid-specific genes all increase in expression when *MATα2* is deleted from an **a**/α cell. The **a**-and α-specific genes are expressed because the proteins that activate them (Mat**a**2 and Matα1) are normally repressed in an **a**/α cell but are expressed when *MATα2* is deleted. Data from one culture of each genotype is plotted here, and data from replicates grown and prepared in parallel are in Supplementary Fig. 5.
b. Mat**a**1 is required for haploid-specific gene repression in *W. anomalus*. mRNA-seq of wildtype *W. anomalus* **a**/α cells (*MAT****a****/MATα*) compared to **a**/α cells with *MAT****a****1* deleted (*MAT****a****/MATα mat****a****1-Δ*). All genes are labeled as in previous panel, except *MAT****a****1* is shown in red, confirming that it has been deleted.
c. Diagram of the sequence upstream of the *RME1* coding sequence indicating presumptive Mat**a**1-Matα2 (green) and Mcm1 (blue) binding sites.
d. Expression levels of endogenous *RME1* transcript (which serves as a control) and various P_*RME1*_-GFP reporter constructs in *W. anomalus* **a** and **a**/α cells measured by RT-qPCR. As indicated, a variety of different deletions were made in the reporter. The results show that the predicted Mcm1-binding site nearest to the transcription start site is necessary for *RME1* repression in the **a**/α cell. Quantities are mean and SD of two cultures grown and measured in parallel, normalized to expression of the housekeeping gene *TBP1*. The experiment was repeated with independent genetics isolates of each strain and each repeat recapitulated the results (Supplementary Fig. 6).

**Figure 4.**
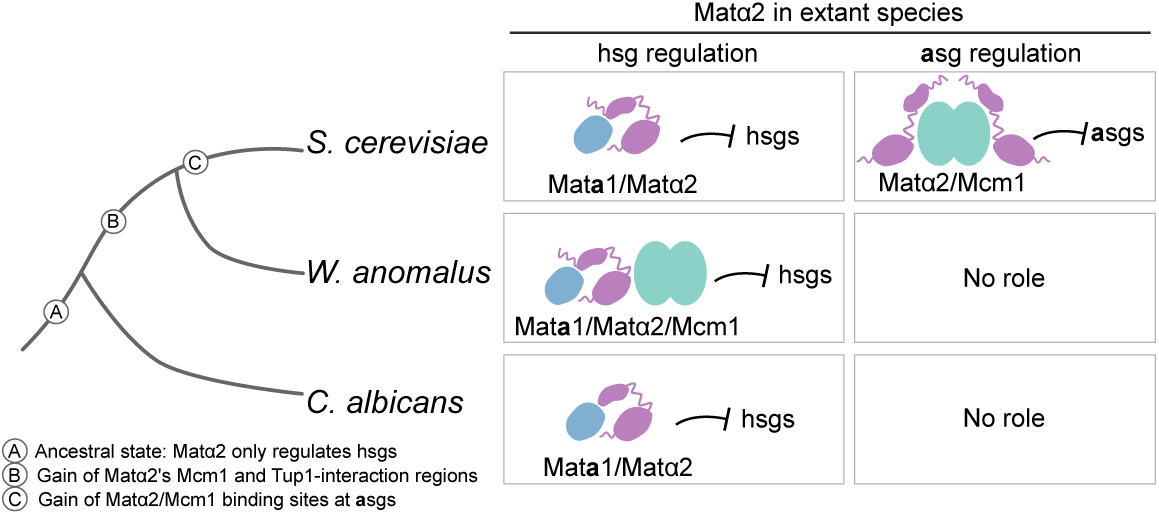
Order of evolutionary events leading to repression of the a-specific genes by Matα2, mapped on to the yeast cladogram. Shown at the right is the role of Matα2 in three extant species. The three-protein solution for repressing the haploid-specific genes is still in place in the *W. anomalus* clade, but in the *S. cerevisiae* lineage was partitioned into **a**-specific gene regulation (which uses only two proteins, Mcm1 and Matα2) and repression of the haploid-specific genes (which requires Matα2 and Mat**a**1). Without the “three-protein” intermediate it is difficult to imagine how **a**-specific gene regulation could have evolved. In particular, the intermediate explains how the necessary changes in the regulatory protein Matα2 could be maintained for millions of years before the new **a**-specific regulatory circuit was completed by acquisition of *cis*-regulatory sequences.

## Discussion

In this paper we examine the evolutionary trajectory through which a new transcriptional circuit was formed in a group of fungal species that includes *S. cerevisiae*. The key regulatory protein responsible for this new circuit (the homeodomain protein Matα2) is ancient, and the genes it controls in the new circuit (the **a**-specific genes) are also ancient. We show that the new, derived circuit—which links Matα2 to the **a**-specific genes by direct binding interactions—formed in at least two stages, separated by millions of years. In the first stage, Matα2 acquired two short amino acid sequences, each of which recognizes an additional ancient protein (Tup1 and Mcm1). In the second stage, the **a**-specific genes acquired *cis*-regulatory sequences recognized by Matα2. The changes in the coding sequence of Matα2 are observed across a broad clade of species, but the *cis*-regulatory changes are seen only in a subset of those species.

This scenario raises the question of how the modifications of Matα2 could arise and be maintained long before the circuit was completed by the *cis*-regulatory changes in the **a**-specific genes. By analyzing a group of species that branched after the modifications in Matα2 had occurred but before the *cis*-regulatory changes were in place, we found the modifications of Matα2 were now needed for Matα2 to carry out its deeply conserved ancestral function. This clade-specific requirement explains how the modifications of Matα2 could be maintained for long periods of evolutionary time before the new **a**-specific gene repression circuit was completed.

This study helps to illuminate several long-standing issues. (1) How is pleiotropy avoided when transcriptional regulators acquire new functions? The modular structure of Matα2 is evident from experiments showing that regions of the protein (Tup1 and Mcm1-interaction regions) that were acquired recently can be transplanted to a variety of outgroup Matα2 proteins (representing the ancestral Matα2) and that they endow the ancestral proteins with the new functions without compromising the existing functions^29^. Matα2 therefore can “cleanly” acquire new functions without compromising existing functions. Virtually all eukaryotic transcriptional regulators have analogous modular structures, and pleiotropy (e.g. off-target effects) can be avoided in the same way. But there is a second, more subtle way that pleiotropy was avoided in the case studied here. The constraints of Matα2’s deeply conserved, ancestral function meant that only certain evolutionary trajectories were available to the protein. Its ancestral function requires of Matα2 to work in combination with Mat**a**1 to bind and repress the haploid-specific genes. Our work shows that there are at least two ways that this binding can occur. It can happen solely through a Mat**a**1-Matα2 heterodimer (as it does the *S. cerevisiae* clade) or it can occur through a triple combination with Mat**a**1, Matα2 and Mcm1 (as is found in *W. anomalus*). All three of these proteins are ancient, and as a default hypothesis, we propose that this shift occurred neutrally with the energy of binding to the control region of the haploid-specific genes simply being “shared out” differently in the two groups of species. According to this idea, the newly acquired interaction between Matα2 and Mcm1 could have occurred only if it was compatible with Matα2’s existing function. In this way, pleiotropy was avoided almost automatically; even before the new **a**-specific gene circuit was formed, the Matα2-Mcm1 combination (which forms the basis of the new circuit) had been “vetted” for millions of years as being compatible with the ancestral functions of Matα2.

(2) Is this evolutionary scenario compatible with the concept of “constructive neutral evolution,” the idea that new functions can evolve through evolutionary transitions of approximately equal fitness^30-32^? Constructive neutral evolution means that new functions need not be dependent on ever-improving performance; rather, complexity can be initially gained neutrally, providing an array of different “solutions” to a regulatory problem. Before the results presented here, it was difficult to understand how the derived circuit represented by *S. cerevisiae* (repression of the **a**-specific genes by Matα2 in α cells) could have evolved because it required changes in both the Matα2 coding region and in the *cis*-regulatory sequences controlling the 5-10 **a**-specific genes. We previously speculated (incorrectly, as it turned out) that the *cis*-regulatory changes in the **a**-specific genes occurred before the coding changes in the Matα2 protein^29^. The present study rules out that hypothesis by showing that the coding-sequence changes must have come first and that they are needed in the extant *W. anomalus* clade for Matα2 to carry out its deeply conserved, ancestral function. We propose that this change represents an example of constructive neutral evolution, in the sense that the neutral sampling of different ways to repress the haploid-specific genes over evolutionary time led to changes in Matα2 that, millions of years later, through exaptation, formed the basis of the new circuit. Without this exploration of different molecular solutions to repress the haploid-specific genes it is difficult to imagine how the new **a**-specific gene circuit could have formed, as it would require numerous coding changes and *cis*-regulatory changes happening more or less simultaneously. Although we cannot rule out the possibility that the changes in the way that the haploid-specific genes were repressed were somehow adaptive, it seems more likely that they occurred neutrally, an explanation consistent with a wide variety of theoretical work^30-33^. In any case, there is no obvious adaptive explanation and neutral evolution is an appropriate default hypothesis.

(3) Is there a “logic” to the structure of transcription circuits? Over the past ten years we have documented many changes in the regulation of the cell-type specific genes (the **a**-specific genes, the α-specific genes, and the haploid-specific genes) across a broad fungal lineage spanning (2) nominally 300 million years. We have observed changes in the DNA-binding specificity of regulatory proteins, the making and breaking of combinations of regulatory proteins, gains and losses of *cis*-regulatory sequences, changes from positive to negative control, changes from direct to indirect regulation, and examples of circuit epistasis. In this paper we show that some clades regulate the haploid-specific genes with a combination of three proteins, while other use only two of the proteins. Throughout all of these changes which have taken place in different lineages, the overall pattern of cell type specific genes expression has largely remained the same: **a**-specific genes are expressed in **a** cells, α-specific genes in α cells, haploid-specific genes in **a** and α cells and all three classes of genes are off in **a**/α cells. If there is any overriding “design logic” to the different mechanisms of regulating these genes, it is difficult to discern. Rather, the spectrum of observed changes seem to reflect a general principle, clearly illustrated by this case: changes in regulatory proteins and *cis*-regulatory sequences that do not compromise existing circuits will be favored. The evolutionary trajectories that are subject to this constraint can provide a source of novelty that can account for the widespread variety in the regulatory mechanisms underlying cell-type specification in fungi. More broadly, the work presented here illustrates that a given transcription circuit is best understood as one of several possible interchangeable solutions rather than as a finished, optimized design^34^.

Their order is addressed in this paper. As indicated in the schematic diagrams, both Matα2 and Mat**a**2 bind DNA with Mcm1, a transcriptional regulator found in all cell types. The schematic diagrams are based on a large body of published literature including genetic, biochemical, and structural studies.

## Supplementary Information

### Yeast cell-type specific gene expression regulation

Saccharomycotina yeast have two mating types termed **a** and α cells that are genetically identical except for a short DNA segment called the mating-type locus (*MAT*)^35^. This segment codes for transcriptional regulators that ensure the proper expression of the **a**, α, and haploid-specific genes. Thus, **a** cells express the **a**-specific and haploid-specific genes (but not the α-specific genes), and α cells express the α-specific and haploid specific (but not the **a**-specific genes) (Fig. 1a). When an **a** cell successfully mates with an α cell, an **a**/α cell is formed. This cell type does not express the **a**, α, or haploid-specific genes but is competent to undergo meiosis. The proper regulation of these three sets of genes, which is directed by the regulatory proteins coded at the mating type locus, is thus essential not only for proper mating, but also for meiosis, which completes the fungal sexual life cycle.

### *W. anomalus* Matα2 has functional Mcm1 and Tup1-interaction regions

Given the similarity of the *W. anomalus* Matα2 protein sequence to that of *S. cerevisiae*, we hypothesized that the *W. anomalus* protein represented the derived form of Matα2. To test this hypothesis, we expressed the *W. anomalus* and, as a control, the *S. cerevisiae MATα2* coding sequences in a *S. cerevisiae* α cell that had the *MATα2* gene deleted and contained a P_*CYC1*_-GFP reporter with an **a**-specific gene *cis*-regulatory sequence taken from the promoter of the *S. cerevisiae* **a**-specific gene *STE2*. As expected, the *S. cerevisiae* Matα2 represses transcription from this promoter by more than 99 percent. In contrast to our expectations, *W. anomalus* Matα2 failed to repress the reporter (Fig. 2b). By constructing a series of chimeric proteins, we mapped the “deficiency” of the *W. anomalus* Matα2 to its homeodomain, indicating that although the protein is in large part conserved, its homeodomain simply did not recognize the *S. cerevisiae* **a**-specific gene *cis*-regulatory sequence (Fig. 2b). To test this idea, we swapped the *S. cerevisiae* homeodomain into *W. anomalus* Matα2 and we found that it could now carry out repression of the reporter construct, although not as well as the bona fide *S. cerevisiae* protein (Supplementary Fig. 1d). *W. ciferrii* and *C. jadinii* Matα2 regions 1 and 2 also resemble the *S. cerevisiae* sequence, and their full-length sequences and the *C. jadinii* homeodomain also do not support **a**-specific gene repression (Supplementary Fig. 1d)

### *W. anomalus* a-specific genes are not directly regulated by Matα2

To determine whether the *W. anomalus* **a**-specific genes are repressed by Matα2, activated Mat**a**2, or both repressed by Matα2 and activated by Mat**a**2, we deleted *MATα2* from an α cell and assayed gene expression changes genome wide by mRNA-seq. None of the **a**-specific genes increased in expression (Fig. 2d, Supplementary Fig. 3), indicating that the *W. anomalus* Matα2 does not directly repress the **a**-specific genes. We further tested this conclusion using a more classical approach. In *S. cerevisiae, MATα2* is necessary for proper mating of α cells; its deletion causes ectopic expression of the **a**-specific genes thereby preventing efficient mating. In *W. anomalus* a wildtype α cell and two independently derived deletions of *MATα2* from α cells all mated with similar efficiency, indicating, unlike in *S. cerevisiae*, that *MATα2* is not required for α cell mating (Fig. 2c). These experiments provide additional support for the conclusion that the *W. anomalus* Matα2 does not regulate the **a**-specific genes in *W. anomalus*.

### *W. anomalus* a-specific genes are activated by Mata2, the ancestral mode of gene expression

The fact that *W. anomalus* Matα2 does not directly repress the **a**-specific genes suggests that, by default, the transcriptional activator Mat**a**2 must positively regulate the **a**-specific genes in *W. anomalus*. To test this hypothesis, we deleted *MAT****a****2* in an **a** cell and measured the expression levels of the **a**-specific genes with the NanoString nCounter system, a method for directly counting individual transcripts of choice in an RNA sample^36^. Each of the **a**-specific genes was significantly under-expressed in the *MAT****a****2* deletion strain with respect to a wildtype **a** cell (Fig. 2e). We also tested the consequences of *MAT****a****2* deletion on **a**-specific mating in a quantitative mating assay and found that an **a** cell with *MAT****a****2* deleted fails to mate at any detectable level in our assay (Fig. 2c). Thus, *W. anomalus* **a**-specific genes are directly regulated by Mat**a**2-activation alone, and Matα2 repression is not involved. Consistent with this conclusion are bioinformatic and experimental analyses of the **a**-specific genes in *W. anomalus, C. jadinii and W. ciferrii*, which identifies *cis*-regulatory sequences that are similar to those controlled by Mat**a**2 in other species (Supplementary Fig. 4a, b). As discussed above, this scheme of regulating the **a**-specific genes is also observed in *C. albicans* and other outgroup species and, by inference, is likely to represent the situation in the last common ancestor of the three clades.

### *W. anomalus* Matα2 retains the deep ancestral function of repressing the haploid-specific genes with Mata**1**

Matα2’s more ancient function is repression of the haploid-specific genes with Mat**a**1. To test whether this ancestral role is retained in *W. anomalus*, we separately deleted either *MATα2* or *MAT****a****1* in an **a**/α cell background and measured gene expression changes by mRNA-seq. In both deletion strains the haploid-specific genes are significantly de-repressed compared to a wildtype **a**/α cell (Fig. 3a, b, Supplementary Fig. 5). We note that deletion of either *MAT****a****1* or *MATα2* also de-represses the *MAT****a****2* and *MATα1* genes, and the **a** and α specific genes are also improperly expressed in the deletion strains. This result also explains how the **a** and α-specific genes are kept off in the **a**/α cells. From these results, we conclude that *W. anomalus* retains the deep ancestral function of Matα2, to repress the haploid-specific genes in combination with Mat**a**1.

### Mcm1 is necessary for haploid-specific gene repression in *W. anomalus*

We introduced a series of *MATα2* constructs into an **a** cell and measured changes in haploid-specific gene expression using the NanoString nCounter system. As expected, addition of a wildtype allele of *MATα2* resulted in transcriptional repression of the haploid-specific genes relative to their expression levels in an **a** cell (Fig. 2c). *RME1*, a deeply conserved haploid-specific gene, was especially tightly repressed, to a level less than one percent of that in **a** cells. Addition of a *MATα2* allele with a point mutation in the Mat**a**1-binding region (region 5), on the other hand, failed to repress *RME1* and the other haploid-specific genes. Addition of *MATα2* chimeras where the Tup1 or Mcm1 interaction regions of Matα2 were replaced by the homologous region of the *C. albicans* (outgroup) Matα2 resulted in substantial loss of repression of many haploid-specific genes. We conclude that, in contrast to the case in *S. cerevisiae*, both the Tup1 and Mcm1-interacting regions of Matα2 are necessary for the complete repression of haploid-specific genes in *W. anomalus.*

The requirement for Matα2 to carry the Mcm1-interaction region in order to repress the haploid-specific genes in *W. anomalus* was surprising because this region is dispensable for haploid-specific gene repression in *S. cerevisiae* and *C. albicans*^29^. We therefore carried out an experiment to test whether Mcm1 has a direct role in repressing the haploid-specific genes in *W. anomalus*. Mcm1 controls the expression of hundreds of genes and is an essential gene in all fungal species tested to date^15^. Therefore, experiments that alter levels of Mcm1 are difficult to rigorously interpret. Instead, we turned to the regulatory region of the *RME1* gene, the most tightly regulated haploid-specific gene, and found two high-scoring matches to the Mcm1 recognition sequence (a *cis*-regulatory sequence that is virtually invariant across these species) (Fig. 3d, Supplementary Fig. 6)^15^. We confirmed that both Mat**a**1 and Matα2 bind at the *RME1* locus by chromatin-immunoprecipitation followed by qPCR (Supplementary Fig. 6b). To test whether Mcm1 is also required for *RME1* repression in an **a**/α cell we constructed a reporter driven by the DNA upstream of the *RME1* gene, showed that it was repressed in **a**/α cells, and tested the effect of deleting the Mcm1 recognition sequences. This experiment has the advantage of maintaining proper regulation of all the normal haploid specific genes, including the endogenous copies of *RME1*.

Deletion of the proximal, but not the distal Mcm1 recognition sequence did away with repression of the *RME1* reporter in **a**/α cells (Fig. 3e, Supplementary Fig. 6c). Based on motif searching, we also identified several potential Mat**a**1-Matα2 binding sequences in the *RME1* control region (Supplementary Fig. 6a). While the two best matches to the Mat**a**1-Matα2 binding site were not required for *RME1* repression, deletion of a possible Matα2 site that abuts the critical Mcm1 site also de-represses *RME1* in an **a**/α cell (Supplementary Fig. 6d). From these results we conclude that Mat**a**1, Matα2, and Mcm1 are all required to repress *RME1* transcription in **a**/α cells. Also required are the regions of Matα2 that bind Mcm1 and Mat**a**1.

## Materials and Methods

### Growth conditions and media

All strains were grown on yeast extract peptone dextrose media (YEPD) at 30°C unless otherwise noted.

### Identification of the Phaffomycetaceae as candidate species

The Phaffomycetaceae form a monophyletic group that branches outside of the last common ancestor of the Saccharomycetaceae (the group that contains the *Saccharomyces* and *Kluyveromyces* species, also known as the “*S. cerevisiae* clade”), as well as the Saccharomycodaceae (an outgroup to the Saccharomycetaceae that includes *Hanseniaspora valbyensis* and other *Hanseniaspora* species). This clade consists of four species that had sequenced genomes at the time this study was conducted: *W. anomalus, W. ciferrii, C. jadinii*, and *C. fabianii*^16,17^. Multiple phylogenomic analyses have placed the Phaffomycetaceae as branching outside of the Saccharomycetaceae and the Saccharomycodaceae with high confidence^16,17^. Other Phaffomycetaceae species lacked sequenced genomes during the time this study was completed, and were therefore excluded from this study. In this study we use the terms “*W. anomalus* clade” and Phaffomycetaceae interchangeably.

### Identification of Phaffomycetaceae mating type transcriptional regulators

The genome sequenced *W. anomalus* strain (NRRL Y-366-8) is an **a** cell. The **a**-mating type locus (*MAT****a***), which encodes the **a**-mating type transcriptional regulators *MAT**a1*** and *MAT****a****2*, is located between the *DIC1* and *SLA2* genes, which are syntenic with the mating type locus in many Saccharomycotina species^17,43^. Unlike *S. cerevisiae, W. anomalus* is not known to undergo mating-type switching, and there is no evidence for a silenced mating-type locus or HO endonuclease in its genome^18,43^. To determine the sequence of the α-mating type locus (*MATα*), we designed oligonucleotides to PCR amplify the region between the *DIC1* and *SLA2* coding sequences of NRRL Y-2153-4, another *W. anomalus* isolate that has been reported to mate with the sequenced **a** cell. This sequencing revealed the genes for the α-mating type transcriptional regulators *MATα1* and *MATα2*. All of the *W. anomalus* mating transcriptional regulator genes except for *MATα1* contain introns, so we amplified and sequenced *MATα2, MAT****a****1* and *MAT****a****2* from cDNA to determine their exon coding sequences. Orthology of the *MAT* genes was further confirmed by best reciprocal BLAST to other Saccharomycotina sequences.

The genome sequenced strain of *W. ciferrii* (NRRL Y-1031) is also an **a** cell, so we used the same approach to determine the sequence of the *W. ciferrii MATα* locus. We amplified and sequenced the region between *W. ciferrii DIC1* and *SLA2* in the complementary mating types NRRL# Y-1031-11 and NRRL# Y-1031-27, and found them to be an **a** cell and an α cell, respectively. The *C. fabianii* genome sequenced strain, NRRL Y-1871, is an α cell, so we were able to directly determine the sequence of *MATα* from the existing sequenced genome. The sequenced *C. jadinii* isolate is homothallic and tetraploid, and the genome therefore encodes both *MAT****a*** and *MATα*. All four Phaffomycetaceae *MATα2* genes contain introns, so for each gene we sequenced the cDNA transcript in order to determine its exonic coding sequence.

### Identification of cell-type specific genes

Phaffomycetaceae haploid, **a**, and *α*-specific genes were identified by best reciprocal TBLASTN to other Saccharomycotina cell-type specific genes, and with the Yeast Genome Analysis Pipeline (YGAP) ^44,45^. In *W. anomalus* the expression patterns of the cell-type specific genes were confirmed by measuring transcript abundance using the NanoString nCounter sytem (method described below) (Supplementary Fig. 2).

### Strain construction in *W. anomalus*

The genome-sequenced strain of *W. anomalus*, NRRL Y-366-8, is also the *W. anomalus* type strain and an **a** cell. To generate an **a**/α strain of *W. anomalus*, NRRL Y-366-8 was plated on media containing 5-FOA to select for spontaneous uracil auxotrophs. An **a** cell auxotrophic for uracil was then crossed to NRRL Y-2153-4, an α cell with a naturally occurring arginine auxotrophy. The resulting **a**/α cell (yCSB 8) was selected for by its ability to grow on synthetic defined media with both arginine and uracil dropped out. The 2N diploid ploidy of yCSB 8 was confirmed by staining with SYTO™ 13 Green Fluorescent Nucleic Acid Stain followed by flow cytometry, and the presence of both *MAT****a*** and *MATα* mating type loci in yCSB 8 was confirmed by PCR.

*W. anomalus MAT****a*** gene deletions were made in NRRL Y-366-8, and MAT**a**/α deletions were made in yCSB 8. We were unable to efficiently transform the NRRL Y-2153-4 α cell strain. In order to generate a more genetically tractable α cell, we sporulated yCSB 8 by growing it on a sterilized carrot slice at room temperature for at least five days^18^. Carrots were peeled, cut into cylindrical wedges, then placed in glass tubes with 1 ml water and autoclaved at 115°C for 15 minutes. 500 μl of cells from liquid culture were collected, washed in water, then resuspended in 200 μl water, layered on top of a sterilized carrot wedge in a six well plate, and placed at room temperature. Spores were visible after five days, and sporulation can be started either from saturated overnight cultures, or mid-log cultures. Spores were then resuspended in 100 μl of 1 mg/ml zymolyase for 11 minutes at 37°C degrees, then dissected. A resulting α cell that proved to be genetically tractable (yCSB 342) was the parent strain for generating the α cell *MATα2* deletions. In addition, the dissected spores included **a** and α cells with arginine auxotrophies that were used as tester strains in quantitative mating assays (described below).

*W. anomalus* genes were deleted using a nourseothricin-resistance marker and a freeze-thaw transformation protocol optimized for use in *W. ciferrii* by Schorsch *et al*.^46^. To construct deletion cassettes, we cloned homology arms into pCS.ΔLig4, a plasmid generated by Schorsch *et al*. ^46^. pCS.ΔLig4 contains *nat*, the nourseothricin-resistance gene, codon-optimized for expression in *W. ciferrii*, and under control of *W. ciferrii P*_*PDA1*_ and *T*_*TEF1*_. It also contains a region of the *W. ciferrii LIG4* gene flanked by loxP sites, for inserting pCS.ΔLig4 into *LIG4* by ends-in recombination. To use the *W. ciferrii-*optimized *nat* marker to delete genes in *W. anomalus* we excised the *LIG4* fragment from pCS.ΔLig4 and ligated in 5’ homology regions in between the SexAI and SalI restriction sites, and 3’ homology regions between the SacI and SpeI restriction sites. Homology regions ranged from approximately 700 basepairs to 2000 basepairs in length. Transformed cells were selected on 50, 200, and 400 μg/ml nourseothricin for NRRL Y-366-8, yCSB 342, and yCSB 8, respectively. Deletions were screened and confirmed by PCR testing for the absence of the deleted ORF, as well as the presence of the junctions between the deletion cassette and the surrounding genome. Approximately 1/500 transformed colonies contained gene deletions, because *W. anomalus*, like many other fungi, seems to favor taking up DNA by non-homologous end joining rather than homologous recombination^46^.

pCS.ΔLig4 also served as the backbone for *MATα2* add-in experiments. A full-length wildtype *MATα2* gene was PCR amplified and inserted into pCS.ΔLig4 between the SexAI and SalI restriction sites, creating pCSB 147. This plasmid was subsequently modified to produce the different alleles of *MATα2* tested in Supplementary Fig. 6 by digesting at either the PstI/XbaI restriction sites or the StyI/BsrGI restriction sites within the *MATα2* coding sequence and ligating in gBlock gene fragments (Integrated DNA Technologies) with specific mutations. These constructs were then linearized by PvuI digestion, transformed into yCSB 5, and allowed to randomly integrate into the genome. At least two independent integrants of each allele was assayed in any experiment involving a randomly integrated construct in *W. anomalus* in order to control for variability due to integration at different loci.

pCSB 147 was modified for chromatin immunoprecipitation of Matα2 by using a gBlock to introduce a 3x HA tag at the N-terminus of *MATα2* at the PstI/XbaI restriction sites. Similarly, to tag *MAT****a****1* we inserted a full length *MAT****a****1* construct with its promoter and terminator into pCS.ΔLig4 at the SexAI/SalI restriction sites. We then introduced a 3x HA tag at the N-terminus of *MAT****a****1* by inserting a gBlock between the XbaI/NcoI restriction sites. Tagged *HA-MATα2* and *HA*-*MAT****a****1* constructs were inserted into *W. anomalus* by random integration.

We also used pCS.ΔLig4 as the basis for a hygromycin-resistance marker optimized for expression in *W. anomalus*. *W. ciferrii* P_*PDA1*_ and T_*TEF1*_ were amplified from pCS.ΔLig4 and cloned into pUC19 at the BamHI/XbaI and PstI/SphI restriction sites, respectively, to create pCSB 137. The coding sequence of hph (a hygromycin-resistance gene) was then codon-optimized for expression in *W. anomalus* using a codon usage table generated by the Kazusa DNA Research Institute (http://www.kazusa.or.jp/codon/), synthesized as a gBlock (IDT), and cloned into pCSB 137 at the XbaI/PstI restriction sites to create pCSB 182. pCSB 182 was used as the backbone for all P_*RME1*_-GFP experiments. First, a GFP coding sequence was also codon-optimized for expression in *W. anomalus*, synthesized as a gBlock with the *W. anomalus ACT1* terminator, and cloned into pCSB 182 at the BamHI/SalI restriction sites. 944 basepairs of the *RME1* regulatory region (immediately upstream of the beginning of the *RME1* ORF) were cloned upstream of GFP by Gibson assembly. Modifications of P_*RME1*_ were subsequently made by inserting gBlocks between the DraIII/SpeI restriction sites that naturally occurred within P_*RME1*_. All P_*RME1*_-GFP constructs were linearized by digestion with SphI and allowed to randomly integrate into the genome of a *W. anomalus* **a** cell (NRRL Y-366-8). Transformants were selected on YEPD plates with 300 μg/ml hygromycin.

### Strain construction in *S. cerevisiae*

All *S. cerevisiae* experiments were performed in the W303 strain, and transformations were performed by a published lithium acetate/single-stranded carrier DNA/polyethylene glycol method^47^. *MATα2* was deleted from the W303 α cell using a KanMX marker amplified with homology to *MATα2.*

*MATα2* expression constructs for **a**-specific gene transcriptional repression assays were cloned into pNH 604, a plasmid that integrates into the *S. cerevisiae* genome in single copy at the *trp1* locus using a *TRP1* marker^29,48^. pNH 604 was digested with SacII and KpnI to excise the Tef overexpression promoter. *MATα2*constructs were amplified from cDNA or synthesized as gBlock synthetic gene fragments (Integrated DNA Technologies), and fused to the *S. cerevisiae MATα2*promoter by fusion PCR, then inserted into pNH 604 between the SacII/KpnI restriction sites. pNH 604 were linearized by PmeI digestion, transformed into the *MATα matα2-Δ* strain, and selected for on SD-trp plates.

The P_*CYC1*_-GFP reporter was designed to integrate in single copy at the *ura3* locus using the hph hygromycin resistance marker. It was adapted from the GFP reporter pTS 61 and the beta-galactosidase reporter pLG699z^11,49^. P_*CYC1*_ was inserted upstream of GFP, with the addition of a KpnI restriction site, allowing *cis*-regulatory elements to be inserted between the XhoI and KpnI sites in P_*CYC1*_, between the upstream activating sequence and transcription start site of the promoter. The hph marker is up stream of P_*CYC1*_-GFP, and the two together are flanked by homology to *URA3*. Overlapping oligos with the **a**-specific gene *cis*-regulatory elements and phosphorylated sticky ends were designed, annealed, and ligated into the XhoI/KpnI sites.

### Design of *MATα2* mutant alleles

To construct chimeric alleles of *MATα2*, we defined the five regions of the Matα2 protein based on the *S. cerevisiae* sequence as previously described (region 1: amino acid 1-21, region 2: amino acid 22-108, region 3: amino acid 109-127, region 4: amino acid 128-188, region 5: amino acid 189-210) ^29^. To introduce a loss of function mutation into region 5 of *W. anomalus MATα2*, we introduced a L215A point mutation^38^

### Gene expression profiling by mRNA-seq

For gene expression profiling by RNA-seq, single colonies were grown to saturation overnight in YEPD, diluted to OD_600_ = 0.1 in fresh YEPD, and then grown for 4-4.5 hours at 30°C. Cells were pelleted by centrifugation at 3000 rpm for 10 minutes, then flash frozen in liquid nitrogen and stored at −80°C. RNA was extracted, purified, and DNase-treated using an Ambion RiboPure RNA Purification kit for yeast. Quality of total RNA was evaluated on an Agilent Bioanalyzer using an Agilent RNA 6000 Pico Kit. Two rounds of polyA selection were performed using the Qiagen Oligotex mRNA Mini Kit, and mRNA quality was evaluated on an Agilent Bioanalyzer using an Agilent RNA 6000 Pico Kit. mRNA samples were then concentrated using a Zymo RNA Clean and Concentrator Kit. cDNA synthesis and library preparation were performed using a NEBNext Directional RNA-seq kit. Libraries were sequenced using single end 50 basepair reads on the Illumina HiSeq 4000 sequencing system in the UCSF Center for Advanced Technologies. Reads were pseudoaligned to all *W. anomalus* coding sequences using kallisto, and differential expression analyses were performed using sleuth^50,51^. Coding sequences were downloaded from the Joint Genome Institute annotation of genes for *Wickerhamomyces anomalus* NRRL Y-366-8 v1.0 ^17^. For differential expression analyses, we focused on the expression patterns of candidate haploid, **a**, and α-specific genes, except in the case of *matα2-Δ* in α cells, where we used a false discovery rate cut off of q < 0.05 to search for differentially expressed genes.

### Gene expression profiling with NanoString nCounter

To measure transcript abundance with the NanoString nCounter system (www.nanostring.com), single colonies were grown to saturation overnight in YEPD, diluted to OD_600_ = 0.1 in fresh YEPD, and then grown for 4-4.5 hours at 30°C. Cells were pelleted by centrifugation for 10 minutes at 3000 rpm, then flash frozen in liquid nitrogen and stored at −80°. RNA was extracted, purified, and DNase-treated using an Ambion RiboPure RNA Purification kit for yeast. NanoString quantification was performed by the UCSF Center for Advanced Technologies using a set of probes designed by NanoString to target the *W. anomalus* mating-type transcriptional regulators, haploid-specific genes, **a**-specific genes, α-specific genes, and a set of non-differentially expressed reference genes (*GAPDH, TBP1, SPC98 and MRPL49*). The NanoString probe for the **a**-mating pheromone gene (*MFA*) targets at least seven paralogs of this gene in *W. anomalus*, because their transcripts are too small and similar in sequence to distinguish from each other by this method. Transcript counts were normalized across samples to the quantities of the non-differentially expressed reference genes.

### GFP reporter assays

Single colonies were grown to saturation overnight in YEPD, diluted to OD_600_ = 0.1 in SD supplemented with amino acids and uracil, and then grown for 4-4.5 hours at 30°C. GFP fluorescence was measured on a BD LSR II flow cytometer. 10,000 cells were measured per sample in each experiment. Cells were gated to exclude debris, and the mean GFP fluorescence was calculated for each reporter strain sample. Three independent isolates of each reporter strain were measured.

### Quantitative mating assays

Quantitative mating assays were performed based on methods described for other yeast species^52^. Strains of interest (for example, gene deletions and their parent strains) were plated on 5-FOA to select for uracil auxotrophs (as described above), and tester strains of the opposite mating type with arginine were isolated from sporulating the **a**/α strain yCSB 8. Single colonies were grown to saturation overnight in YEPD, diluted to OD_600_ = 0.15 in fresh YEPD, and then grown to mid-log phase and combined in a final volume of 5 ml. 1 ml (about 1×10^7^ cells) of the mating type present in excess was combined with 200 μl of the strain to be measured, for a final ratio of 5:1 strain in excess to limiting strain, by volume. The mating mixes were then deposited onto 0.8 μm nitrocellulose filters using a Millipore 1225 Vacuum Sampling Manifold. The filters were then placed on a YEPD agar plate and incubated for 5 hours at 30°C. The cells were resuspended by placing the filters in 5 ml water, and cells dispersed by vortexing. Dilutions were plated on SD-arg-ura plates to select for conjugants, and SD-ura to select for the limiting parent as well as to conjugants. After 2-3 days of growth at 30°C, colonies were counted, and the mating efficiency was calculated by the following formula: Mating efficiency = (mating products) / (mating products + limiting tested parents) × 100%. In other words, Mating efficiency = (number of colonies on SD-arg-ura)/(number of colonies on SD-ura) × 100%.

### *Cis*-regulatory motif discovery and searching

A *cis*-regulatory motif for the Phaffomycetaceae **a**-specific genes was generated using MEME^41^. We used sequences 500 basepairs upstream of the **a**-specific gene orthologs in *W. anomalus* and *W. ciferrii* and used MEME to search for motifs between 24 and 27 bases wide occurring zero or one time in those sequences. These sequences were from the *W. ciferrii STE2, STE6*, and *AXL1* genes, and the *W. anomalus STE2, STE6, ASG7, AXL1, BAR1*, and *STE14* genes. This returned a motif that resembled a Mat**a**2-Mcm1 binding site. We used the TOMTOM tool in the MEME suite to align and compare this *W. anomalus*/*W. ciferrii* **a**-specific gene motif to both the *S. cerevisiae* Matα2-Mcm1 motif and the *C. albicans* Mat**a**2-Mcm1 motif^40^. We searched for Matα2, Mat**a**1, Mat**a**2, and Mcm1 *cis*-regulatory motifs using the MAST and FIMO tools in the MEME suite.

### RT-qPCR

Single colonies were grown to saturation overnight in YEPD, diluted to OD_600_ = 0.1 in fresh YEPD, then grown for 4-4.5 hours at 30°C. Cells were pelleted by centrifugation for 10 minutes at 3000 rpm, then flash frozen in liquid nitrogen and stored at −80°. RNA was purified using the MasterPure™ Yeast RNA Purification Kit from Epicentre. cDNA was synthesized with random hexamer primers using SuperScript**^®^** IV reverse transcriptase. qPCR was performed using BioRad SYBR Green Master Mix on a BioRad CFX Connect™ Real-Time PCR Detection System. Each sample was quantified in three technical replicates, and each experiment was performed with two biological replicates per sample. Relative quantifications were determined using a standard curve, and quantifications for *RME1* and *GFP* were normalized to the non-differentially regulated housekeeping gene *TBP1*.

### Chromatin-immunoprecipitation

Chromatin immunoprecipitation (ChIP) was performed as previously described^11,53^. Briefly, single colonies were grown to saturation overnight in YEPD, diluted to OD_600_ = 0.1 in fresh YEPD, then grown at 30°C to OD_600_ = 0.4. Cells were then crosslinked with formaldehyde, quenched with glycine, washed with TBS (20mMTris/HCl pH 7.4, 150 mM NaCl), flash frozen in liquid nitrogen, and stored at −80°C. Cells were lysed at 4°C in lysis buffer (50 mM HEPES/KOH pH 7.5, 140 mM NaCl, 1mM EDTA, 1% Triton X-100, 0.1% Na-Deoxycholate) plus protease inhibitors (Roche Complete Protease Inhibitor Cocktail EDTA-free) with 0.5 mm glass beads on a vortex for two hours. Recovered chromatin was then sheared via sonication on a Diagenode Biorupter (three ten minute periods of sonication on level 5 for 30 seconds, then off for 1 minute). 10 mg of anti-HA 12CA5 from Roche was used to bind the HA-tagged transcription factors overnight at 4°C in lysis buffer plus protease inhibitors, and protein-G sepharose beads were used to collect immunoprecipitated DNA. Immunoprecipitated DNA was then recovered by washing with lysis buffer, wash buffer (10mM Tris/HCl pH 8.0, 250mM LiCl, 0.5% NP-40, 0.5% Na-Deoxycholate, 1mM EDTA), and TE, then collected in elution buffer (50mM Tris/HCl pH8.0, 10mM EDTA, 1%SDS). Crosslinking was reversed by Proteinase K treatment for two hours at 37°C, then16 hours at 65°C. DNA was then cleaned up using the Qiagen MinElute kit.

*HA-MAT****a****1* and *HA-MATα2* constructs were inserted into the *W. anomalus* genome via random integration. Two to four independent isolates of each strain were processed, and two samples of untagged **a**/α cells, and one each of untagged **a** and untagged α cells were also processed as negative controls. Abundance of the *RME1* promoter region in tagged vs. untagged samples was measured by qPCR and compared to the promoter regions of the housekeeping genes *TBP1, ACT1*, and *RPS20*, and the **a**-specific gene *STE2*.

### Data availability

mRNA-seq data that supports the findings of this study is in the process of being submitted to the NCBI Gene Expression Omnibus.

## Acknowledgements

We thank L. Noiman, M. Lohse, K. Fowler, and L. Booth for comments on the manuscript, and C. Baker, I. Nocedal, C. Dalal, and N. Ziv for technical help. We thank Christoph Schorsch of Evonik Industries for providing us with the plasmid used to genetically modify *W. anomalus*. The work was supported by grant R01 GM037049 from the National Institutes of Health. C.S.B. was supported by an Achievement Rewards for College Scientists (ARCS) Scholarship from the ARCS Foundation. T.R.S. was supported by a Graduate Research Fellowship from the National Science Foundation.

## Author Contributions

C.S.B., T.R.S., and A.D.J. designed and interpreted experiments and edited the manuscript. T.R.S. performed preliminary bioinformatic analyses of Matα2 and **a**-specific gene *cis*-regulatory sequences and sequenced the *W. anomalus* α mating-type locus. C.S.B. performed all experiments and other bioinformatic analyses. C.S.B. and A.D.J wrote the manuscript.

---

**Supplementary Figure 1.**
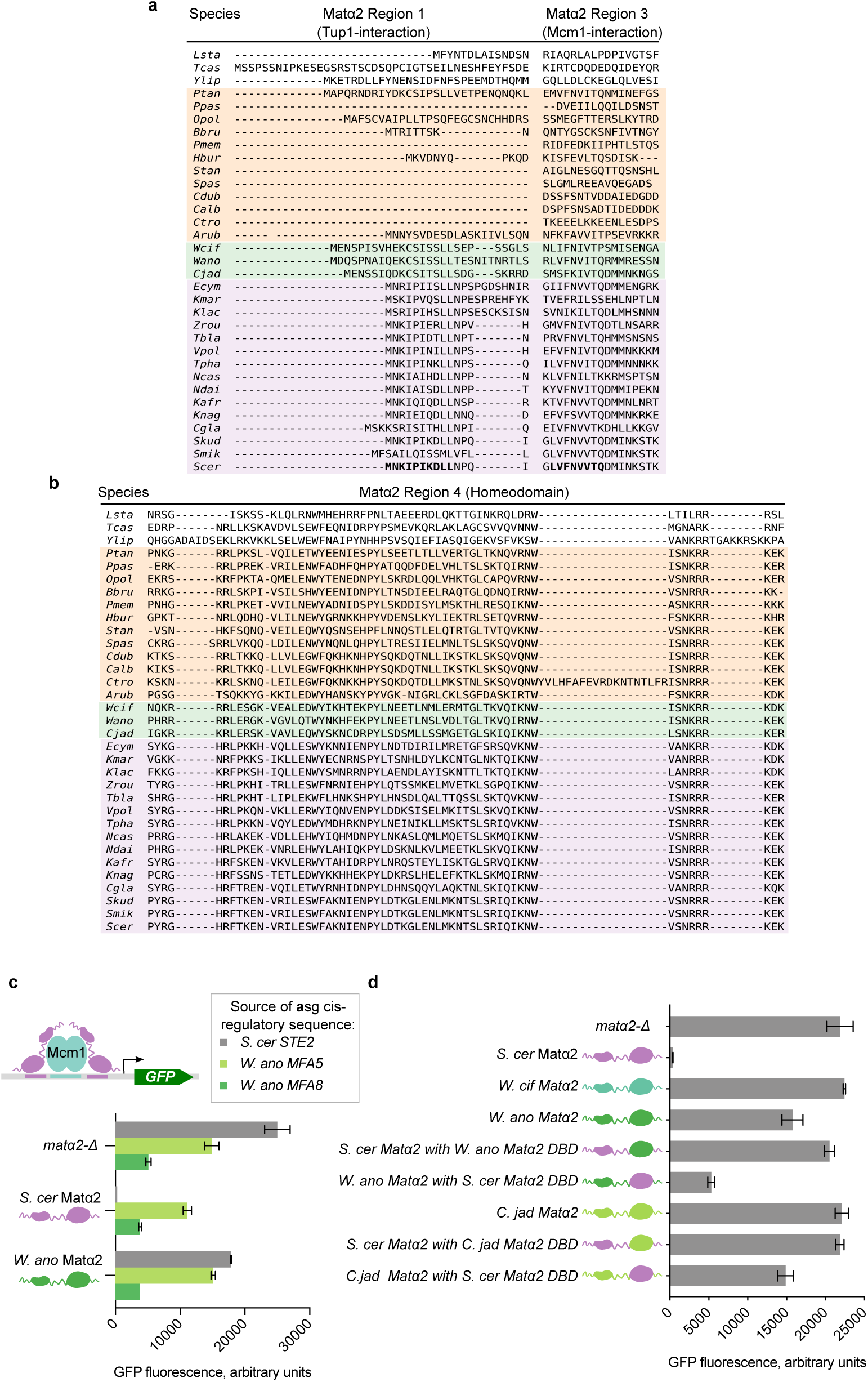
*W. anomalus* clade Matα2 orthologs contain functional Tup1 and Mcm1-interaction regions, but cannot repress a-specific gene transcription due to changes in homeodomain. a. Multiple sequence alignment of Matα2 regions 1 and 3, generated by MUSCLE. *S. cerevisiae* clade (Saccharomycetaceae) sequences are shown in purple, *W. anomalus* clade (Phaffomycetaceae) sequences are shown in green, and *C. albicans* clade (Pichiacea and Debaryomycetaceae) sequences are shown in orange. Species are denoted with a four letter code where the first letter is the first letter of the genus name, and the next three letters are the first three letters of the species name^20,21,37,38^. Residues known to be functionally important for binding Tup1 and Mcm1 in *S. cerevisiae* are bolded on the *Scer* line.
b. Multiple sequence alignment of Matα2 region 4, the homeodomain, generated by MUSCLE. As above, *S. cerevisiae* clade (Saccharomycetaceae) sequences are shown in purple, *W. anomalus* clade (Phaffomycetaceae) sequences are shown in green, and *C. albicans* clade (Pichiacea and Debaryomycetaceae) sequences are shown in orange.
c. The **a**-specific gene *cis*-regulatory elements from the *S. cerevisiae STE2* gene, and two *W. anomalus MFA* genes were placed into a reporter construct and used to test the ability of *S. cerevisiae* Matα2 (purple) and *W. anomalus* Matα2 (green) to repress **a**-specific gene transcription. *MATα2* constructs were driven by the *S. cerevisiae MATα2* promoter and inserted into a *S. cerevisiae* α cell with *MATα2* deleted (*matα2-Δ*). Mean and SD of three independent isolates grown and tested in parallel are shown for all GFP reporter experiments.
d. *W. anomalus* clade Matα2 proteins, as well as chimeric Matα2 alleles with the *S. cerevisiae* Matα2 homeodomain DNA-binding domain (“DBD”) (purple) swapped into *W. anomalus* clade Matα2 protein sequences (*W. anomalus, W. ciferrii*, and *C. jadinii*) (shades of green), and vice versa, were tested for their ability to repress transcription of a reporter with the Matα2-Mcm1 binding site from *S. cerevisiae STE2* promoter. *W. anomalus* clade Matα2s consistently do not support **a**-specific gene transcriptional repression in this assay. For both *W. anomalus* and *C. jadinii*, this is due to the DNA-binding domain. Mean and SD of three independent genetic isolates grown and tested in parallel are shown.

**Supplementary Figure 2.**
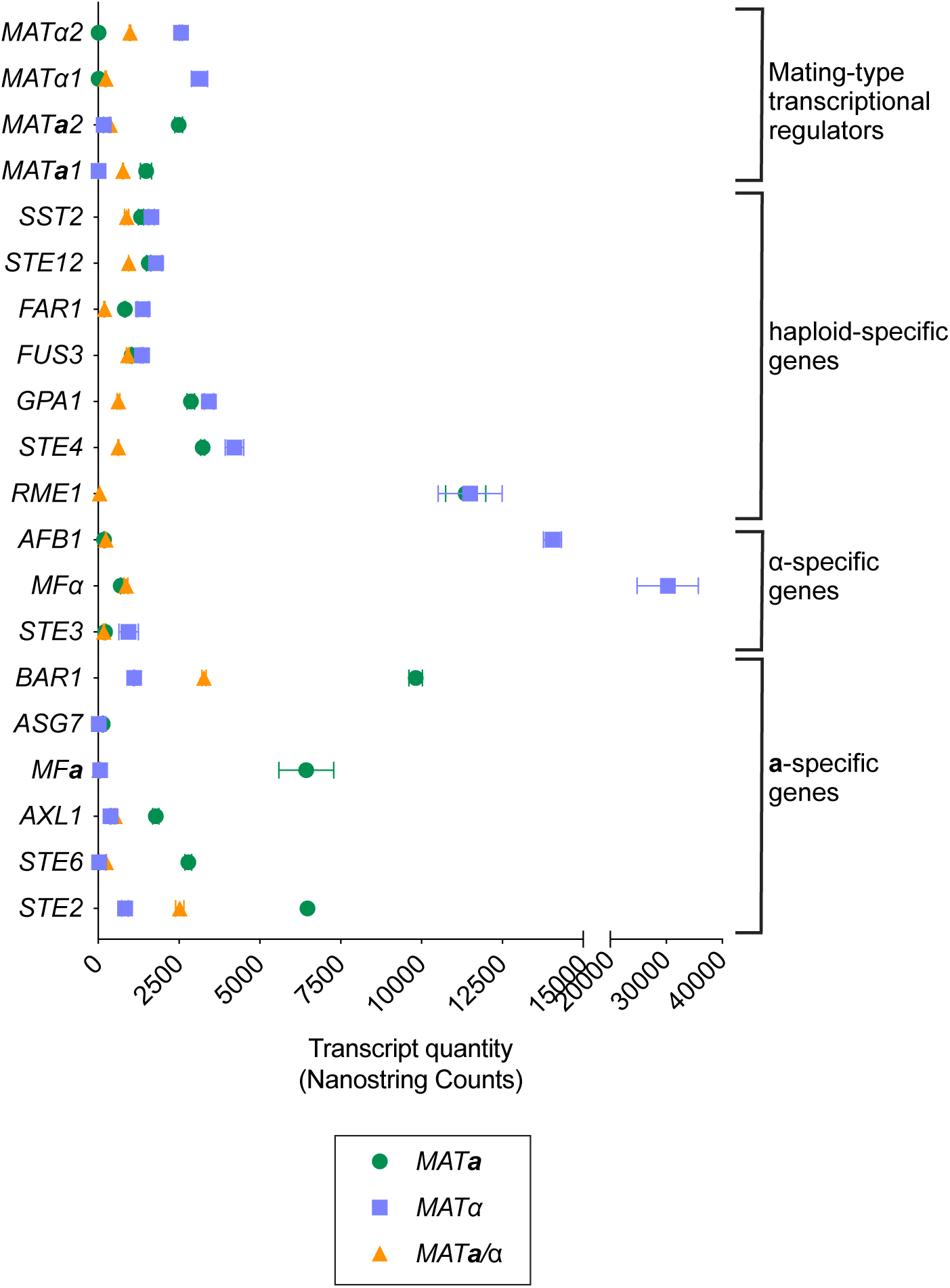
Cell-type specific gene expression patterns in *W. anomalus*. Cell-type specific gene expression levels in wildtype *W. anomalus* **a** (green circles), α (blue squares), and **a**/α (orange triangles) cells measured with the NanoString nCounter system. Mean and SD of two samples are plotted, except for cases in which the error bars are smaller than the data point. This is not intended to be an exhaustive list, and there may be additional cell-type specific genes.

**Supplementary Figure 3.**
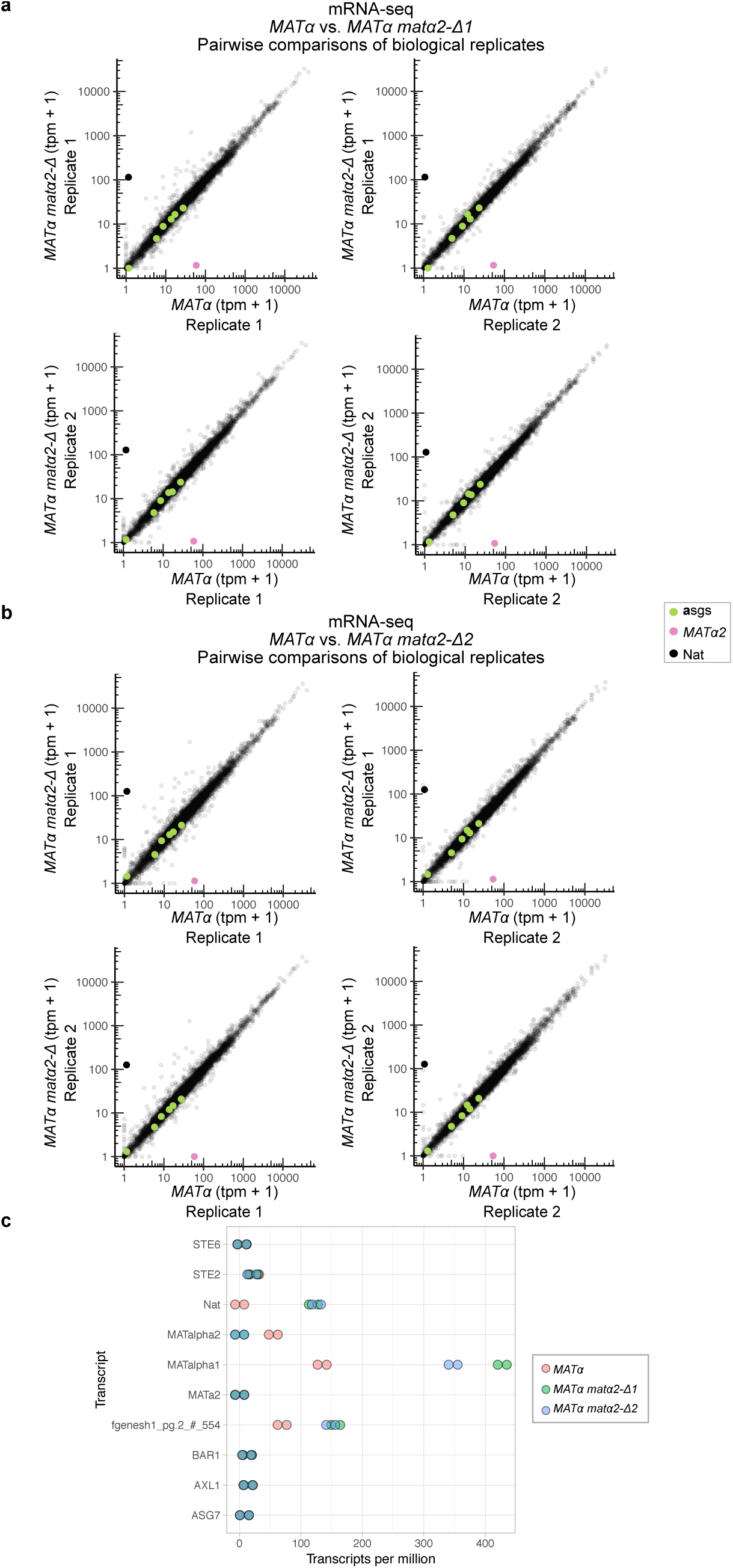
*W. anomalus MATα2* does not directly repress the a-specific genes. a. mRNA-seq of wildtype *W. anomalus* α cells (*MATα*) vs. first isolate of α cells with *MATα2* deleted (*MATα2 matα2-Δ 1*). *MATα2* expression in transcripts per million (tpm) is shown in pink, *Nat* expression is shown in opaque black, the **a**-specific genes *STE2, AXL1, ASG7, BAR1, STE6* and *MAT****a****2* are shown in green, and all other mRNAs are shown in translucent black. Replicate 1 vs. Replicate 1 plot is the same as shown in main text Fig. 2d, b other plot are pairwise comparisons of the other biological replicates performed in parallel in this experiment.
b. mRNA-seq of wildtype *W. anomalus* α cells (*MATα*) vs. second isolate of α cells with *MATα2* deleted (*MATα2 matα2-Δ 2*). *MATα2* expression in transcripts per million (tpm) is shown in pink, *Nat* expression is shown in opaque black, the **a**-specific genes *STE2, AXL1, ASG7, BAR1, STE6*, and *MAT****a****2* are shown in green, and all other mRNAs are shown in translucent black. The wildtype α cell data is the same as in Supplementary Fig. 3a.
c. Expression levels of genes of particular interest from mRNA-seq experiments plotted above, in transcripts per million (tpm). Shown are **a**-specific genes (*STE2, AXL1, ASG7, BAR1*, and *STE6*), mating-type transcriptional regulators (*MAT****a****2, MATα2, and MATα1*), the drug-resistance marker used to delete *MATα2* (Nat), and the transcript “fgenesh1_pg.2_#_554,” which, besides *MATα2, and MATα1* and Nat, is the only significantly differentially regulated transcript shared between both isolates of *MATα2 matα2-Δ vs. MATα2* (at a false discovery rate q<0.05, calculated using the RNA-seq analysis software sleuth).

**Supplementary Figure 4.**
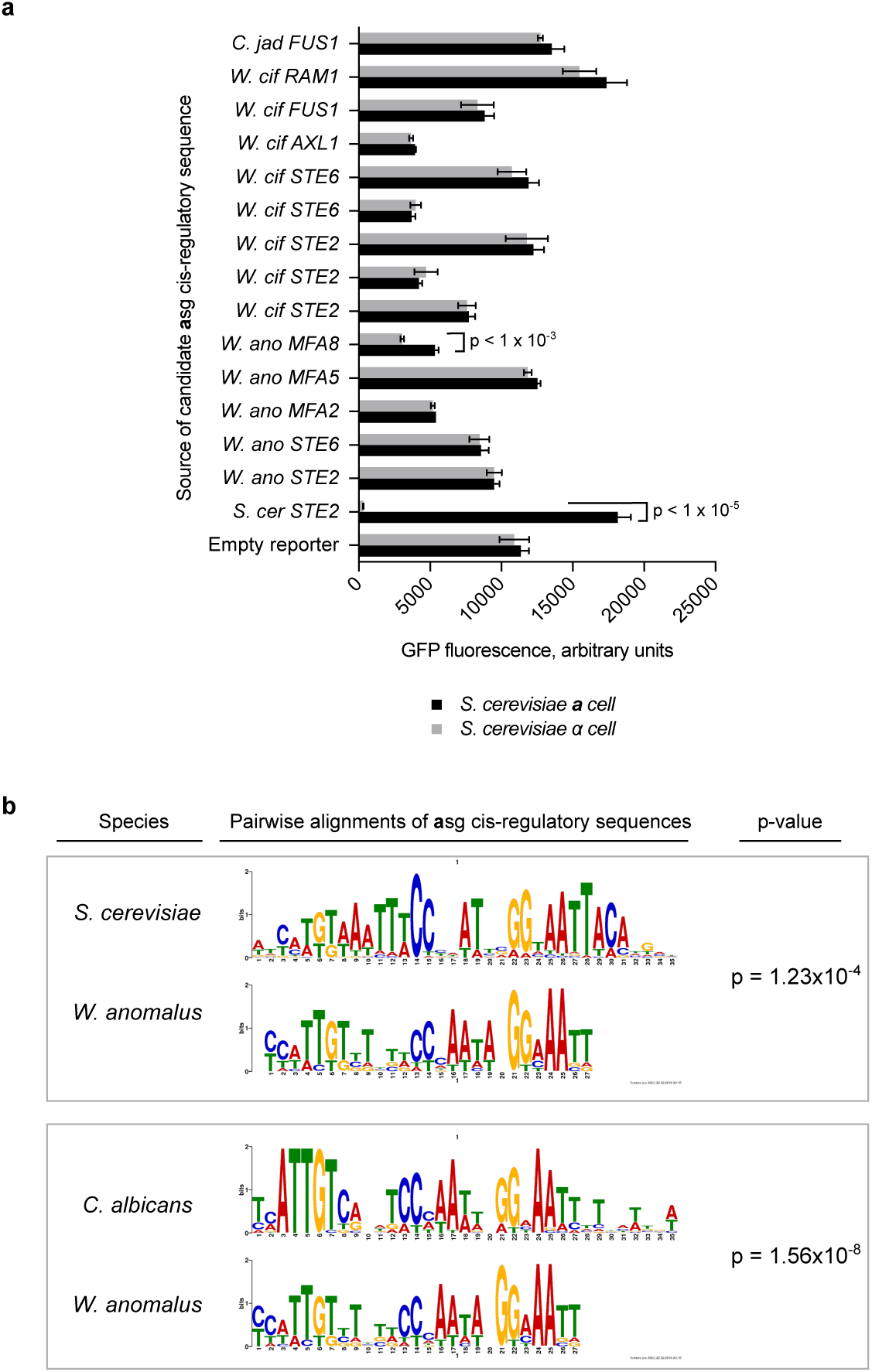
*W. anomalus* a-specific genes are activated by Mata2, the ancestral. a. We identified matches to both the *C. albicans* Mat**a**2-Mcm1 and the *S. cerevisiae* Matα2-Mcm1 binding sites in the promoters of the *W. anomalus, W. ciferrii*, and *C. jadinii* **a** specific genes^39^. These putative *cis*-regulatory sequences were cloned into the *S. cerevisiae* P_*CYC1*_*-*GFP reporter and transformed into *S. cerevisiae* **a** and α cells to test for Matα2-dependent transcriptional repression. While the *S. cerevisiae STE2 cis*-regulatory sequence supports transcriptional repression by Matα2, none of the *W. anomalus* clade **a**-specific gene *cis*-regulatory sequences supported repression by Matα2 to the same extent. Only one of the 14 CREs tested, found upstream of a paralog of the *W. anomalus* **a**-mating pheromone gene *MFA (*“*MFA8*”), showed a statistically significant difference between **a** and α cells with 1.75-fold repression (p < 0.05). This sequence is also the best match to the *S. cerevisiae* Matα2-Mcm1 binding site that we identified in the *W. anomalus* clade. Given the overall evidence for a lack of *W. anomalus* clade Matα2 repression of the **a**-specific genes, we do not believe that this single instance of partial repression by *S. cerevisiae* Matα2 reflects actual or vestigial Matα2-Mcm1 repression in *W. anomalus*.
b. *De novo* motifs for the *W. anomalus* clade **a-**specific genes were generated using the motif-elicitation software MEME, and compared to both the *C. albicans* **a**-specific gene *cis*-regulatory sequence (Mat**a**2-Mcm1 binding site) and the *S. cerevisiae* **a**-specific gene *cis*-regulatory sequence (Matα2-Mcm1 binding site) using Tomtom, a tool within the MEME suite^40,41^. While the *W. anomalus* clade **a**-specific gene *cis*-regulatory sequence shares homology with both the *S. cerevisiae* and *C. albicans* sequences due to the shared Mcm1-binding site, it is more similar to the *C. albicans* site than the *S. cerevisiae* one.

**Supplementary Figure 5.**
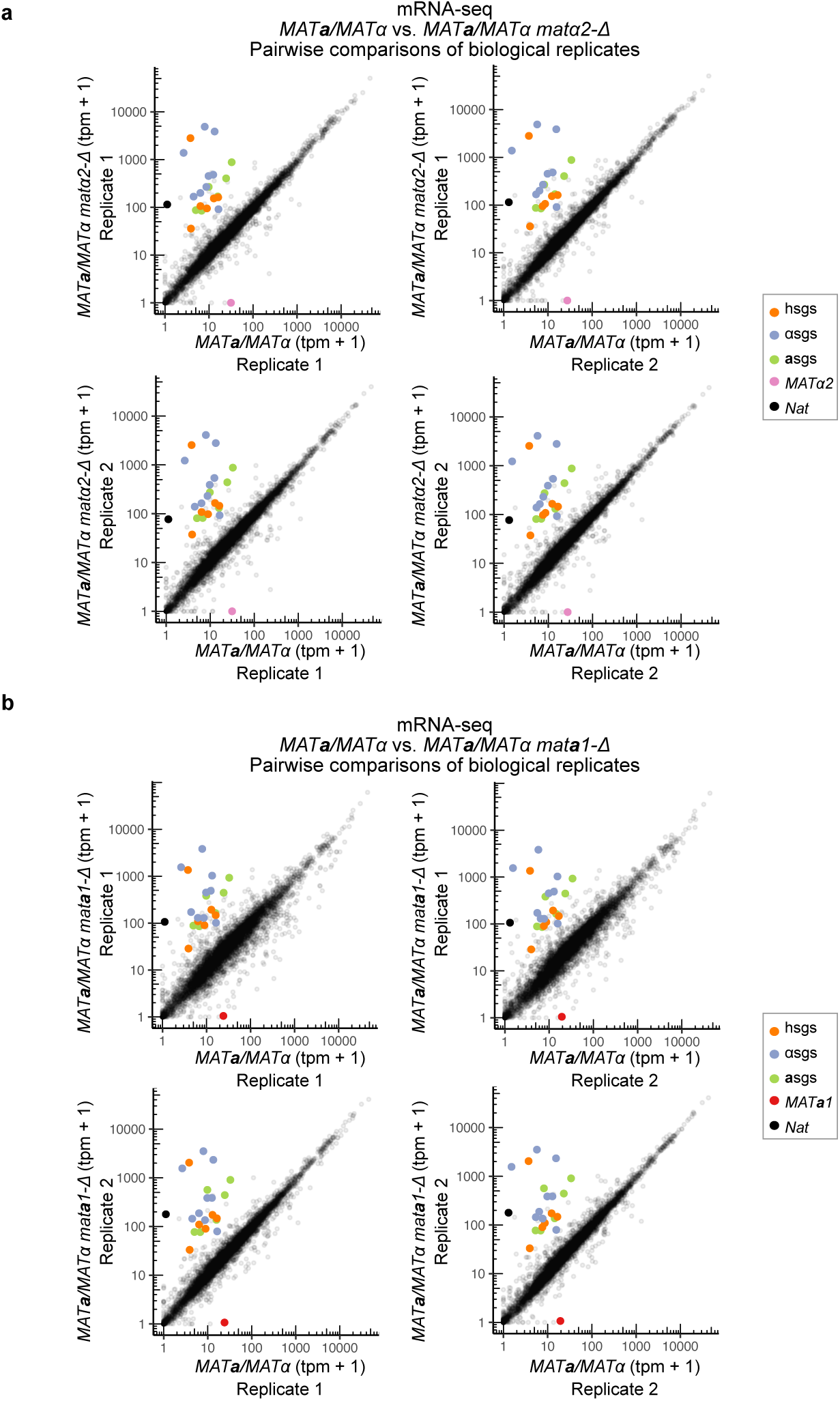
The *W. anomalus* haploid-specific genes are repressed by Matα2 and Mat>a1. a. mRNA-seq of wildtype *W. anomalus* **a**/α cells (*MAT****a****/MATα*) vs. **a**/α cells with *MATα2* deleted (*MAT****a****/MATα matα2-*Δ). *MATα2* expression in transcripts per million (tpm) is hown in pink, the **a**-specific genes *STE2, AXL1, ASG7, BAR1, STE6* and *MAT****a****2* are shown in green, the haploid-specific genes *STE4, GPA1, FUS3, SST2, RME1*, and *FAR1* are shown in orange, the α-specific genes *SAG1, STE3, STE13, MFα* (multiple copies), *AFB1* (multiple copies), and *MATα1* are shown in blue-gray, the selective marker used to delete *MATα2* is shown in opaque black, and all other mRNAs are shown in translucent black. The **a**-, α-and haploid-specific genes are more highly expressed when *MATα2* is deleted. Replicate 1 vs. Replicate 1 plot is the same as shown in main text Fig. 3a, b other plot are pairwise comparisons of the other biological replicates performed in parallel in this experiment.
b. mRNA-seq of wildtype *W. anomalus* **a**/α cells (*MAT****a****/MATα*) vs. **a**/α cells with *MAT****a****1* deleted (*MAT****a****/MATα mat****a****1-Δ*). Genes are highlighted as above, except *MAT****a****1* is shown in red and *MATα2* is not highlighted. As when *MATα2* is deleted from an **a**/α cell, the **a**-, α-and haploid-specific genes are more highly expressed when *MAT****a****1* is deleted. Replicate 1 vs. Replicate 1 plot is the same as shown in main text Fig. 3b, b other plot are pairwise comparisons of the other biological replicates performed in parallel in this experiment.

**Supplementary Figure 6.**
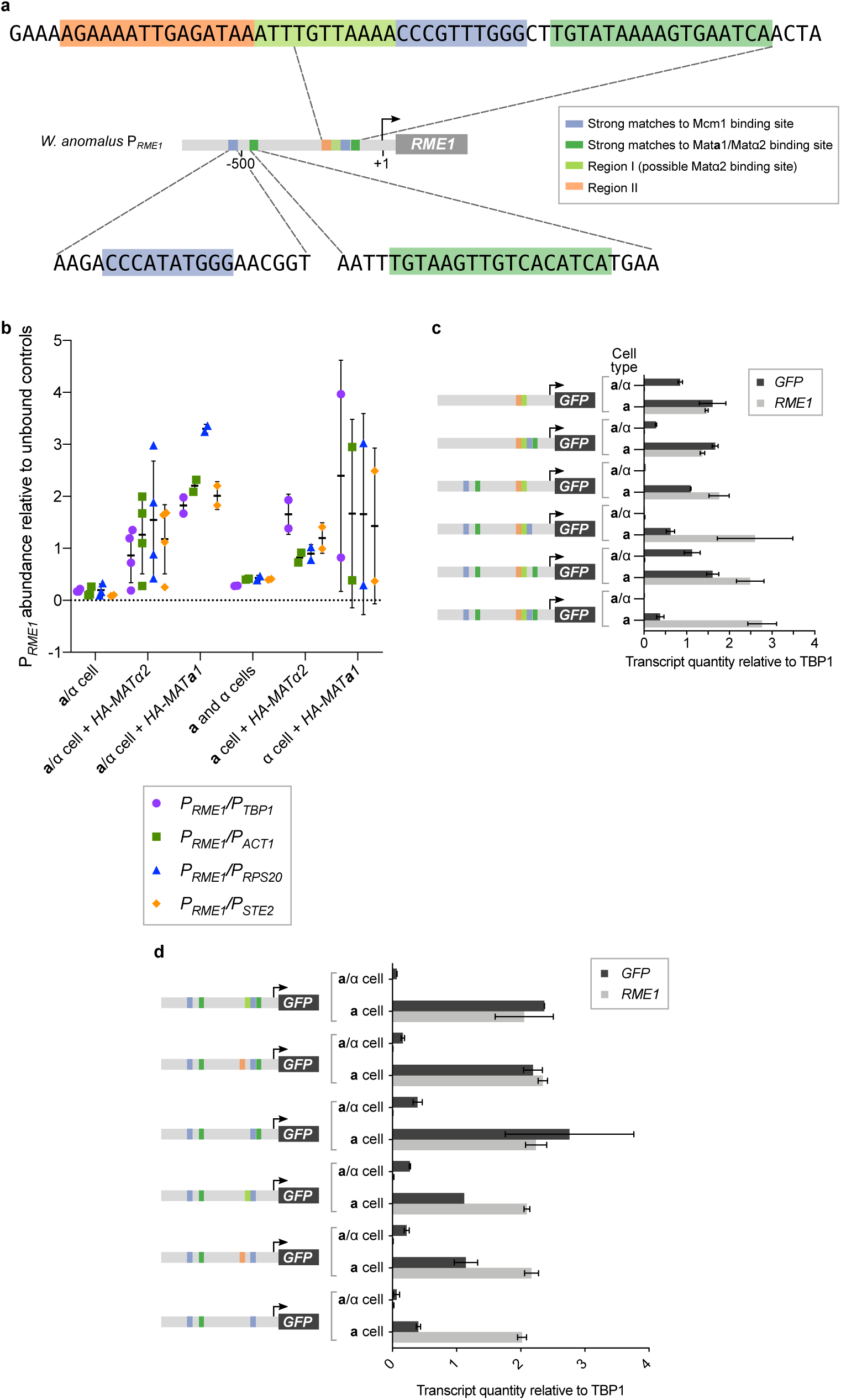
Matα2, Mata1, and an Mcm1 *cis*-regulatory sequence are directly required for *RME1* repression in a/α cell. a. Diagram of the sequence upstream of the *RME1* coding sequence indicating the two strongest matches to Mat**a**1-Matα2 (green) and Mcm1 (blue) binding sites detected, as well as their location relative to the transcription start site. In addition, the smaller lime green box (Region I) indicates a possible Matα2 binding site, and Region II indicates another stretch of sequence to the left of the proximal Mcm1 site that we experimented with. The TGT residues within Region I may be a possible Matα2 binding site and are positioned relative to the adjacent Mcm1 binding site as they would be in a *S. cerevisiae* Matα2-Mcm1 **a**-specific gene *cis*-regulatory site.
b. Matα2 and Mat**a**1 were N-terminally HA-tagged and transformed into *W. anomalus* cells. Chromatin immunoprecipitation (ChIP) using anti-HA antibodies was performed with wildtype *W. anomalus* cells as an untagged control, followed by qPCR to measure abundance of P_*RME1*_ relative to several unbound regions as controls (P_*TBP1*_ in purple circles, P_*ACT1*_ in green squares, P_*RPS20*_ in blue triangles, and P_*STE2*_ in orange diamonds). Results from individual ChIP samples are plotted as dots, and lines show the mean and SD for each strain background.
c. Expression levels of endogenous *RME1* transcript and P_*RME1*_-GFP reporter (*GFP*) in *W. anomalus* **a** and **a**/α cells measured by RT-qPCR. Mat**a**1-Matα2 (green) and/or Mcm1 (blue) binding sites were deleted from P_*RME1*_-GFP individually and in combination, and the reporters were transformed into an **a** cell. Orange and lime green boxes show the Region I and II *cis*-sequences to the left of the proximal Mcm1 start site that are unaltered in this experiment. The **a** cells were crossed to α cells to form **a**/α cells, and expression levels of the reporter and the endogenous transcript were measured in both cell types. The proximal Mcm1-binding site with respect to the transcription start site is necessary for *RME1* repression in the **a**/α cell. Quantities are mean and SD of two cultures grown and measured in parallel, normalized to expression of the housekeeping gene *TBP1*. This is a repeat of the experiment shown in Fig. 3e, b independent genetic isolates of the strains.
d. Expression levels of endogenous *RME1* transcript and P_*RME1*_-GFP reporter (*GFP*) in *W. anomalus* **a** and **a**/α cells measured by RT-qPCR. Possible Mat**a**1-Matα2 binding sites (orange and lime green boxes) immediately to the left the proximal Mcm1 site were deleted from P_*RME1*_-GFP individually and in combination, and the reporters were transformed into an **a** cell. The **a** cells were then crossed to α cells to form **a**/α cells, and expression levels of the reporter and the endogenous transcript were measured in both cell types. While the sequences to the right of the proximal Mcm1 site strongly resemble the Mat**a**1-Matα2 binding site from other yeast species, it is actually the sequences to the left of this Mcm1 *cis*-regulatory site that are necessary for complete *RME1* repression in the **a**/α cell^12^. Quantities are mean and SD of two cultures grown and measured in parallel, normalized to expression of the housekeeping gene *TBP1*.

